# Single-cell dynamics of core pluripotency factors in human pluripotent stem cells

**DOI:** 10.1101/2022.12.13.520282

**Authors:** Sonja Mihailovic, Samuel C. Wolff, Katarzyna M. Kedziora, Nicole M. Smiddy, Margaret A. Redick, Yuli Wang, Guang Ken Lin, Tarek M. Zikry, Jeremy Simon, Travis Ptacek, Nancy L. Allbritton, Adriana S. Beltran, Jeremy E. Purvis

## Abstract

The human transcription factors OCT4, SOX2, and NANOG form a core signaling network critical for maintaining stem cell pluripotency and self-renewal potential. The spatiotemporal expression dynamics of these pluripotency factors throughout differentiation is unclear, limiting our understanding of stem cell fate decisions. Here, we combined CRISPR/Cas9-mediated gene editing with microraft array technology to generate human embryonic stem cell lines with endogenously tagged fluorophores for OCT4, SOX2, and NANOG. Fluorescence time-lapse imaging revealed that pluripotent stem cells show gastrulation-like patterning without direct chemical induction. Directed differentiation to the three primary germ layers—endoderm, mesoderm, and ectoderm—revealed distinct spatiotemporal patterns of SOX2 and NANOG expression in single cells. Finally, we captured dynamic changes in cell morphology during ectoderm differentiation corresponding to the formation of neural rosettes. This study provides a robust method for generating live-cell reporters in human stem cells and describes the single-cell dynamics of human pluripotency factors during differentiation.

## Introduction

The ability to study human embryonic stem cells (hESCs) from previously derived and established cell lines has accelerated our knowledge of how human pluripotent stem cells maintain pluripotency and differentiation capabilities^1–4^. These cells are derived from the inner cell mass of human embryos and have the unique cellular features of self-renewal and the capacity to differentiate into virtually any human cell type^5–7^.

Human ESCs maintain their pluripotency through a signaling network involving the core transcription factors, OCT4, SOX2, and NANOG^8,9^. In general, this signaling network maintains pluripotency by binding to and promoting gene expression of genes controlling self-renewal and pluripotency while inhibiting genes associated with differentiation^10,11^. These three transcriptional regulators also bind to their own and each other’s promoters, creating a positive feedback loop that reinforces expression of all three gene products^12–14^.

Perturbations to this stable self-reinforcing network leads to differentiation toward one of the three primary germ layers: ectoderm, mesoderm, or endoderm^15–17^. In vivo, these differentiation events occur in a coordinated, self-organizing process called gastrulation, which occurs ~3 weeks post-fertilization for human embryos^18^. As individual hESCs undergo differentiation, each germ layer exhibits a precise expression pattern for OCT4, SOX2, and NANOG that determines lineage specification^19–21^.

For example, during endoderm cell fate decisions, SOX2 is silenced first while NANOG expression is prolonged, allowing NANOG to form a complex with SMAD2/3^22–24^. This complex then interacts and binds with Eomesodermin (EOMES), a T-box transcription factor involved with endoderm specification, and acts as an activator for additional endoderm genes^25^. As NANOG expression is lost, EOMES and SMAD2/3 interact to continue the specification to the endoderm lineage^23,26^.

In mesoderm, expression of SOX2 is sustained longer than in definitive endoderm as PRMT8 prolongs SOX2 expression to promote AKT signaling^27^. Then, SOX2 expression is repressed by the MSX2 homeobox transcriptional regulator, which activates the BMP pathway to induce NODAL expression and differentiation into mesendoderm tissue^28^. Additionally, the Brachyury (BRA) transcription factor is also expressed during mesendoderm formation but is required for mesoderm specification as BRA promotes essential mesoderm genes such as WNT-3A and TBX6 while additional NANOG suppression induces the formation of specified mesoderm subtypes^22,29^.

During ectoderm differentiation, SOX2 is consistently expressed and eventually increases expression as cells differentiate along the neuroectoderm lineage to neural progenitor cells (NPCs)^12,15,30,31^. This increase in SOX2 expression allows SOX2 to bind to and form a complex with PAX6 to induce neuroectoderm and NPC cell fate renewal pathways^32,33^. This pattern of sustained SOX2 expression contrasts mesoderm and definitive endoderm, where the lack of SOX2 acts as a repressor for the neuronal lineage^34^. Thus, the temporal expression dynamics for OCT4, SOX2, and NANOG follow an intricate pattern that aid in guiding the developmental fate decisions of individual stem cells^9,11,20^.

Much of our knowledge of these expression patterns comes from *in vitro* models that explore the developmental processes involved in the cell fate decisions made during differentiation and embryonic development. These models range from separated differentiations down a specified germ layer, 2D self-organizing models such as the micropattern-based gastruloids, and 3D gastruloids models such as the free-floating gastruloid^35–37^. These systems typically utilize fixed cells because the development of live cell reporters to monitor real-time expression^38–42^ is challenging. The genetic manipulation required for these reporters is known to potentially trigger chromosome instability, genomic mutations, and unsupervised differentiations^43–45^.

Thus, there is a need to understand the temporal expression dynamics of OCT4, SOX2, and NANOG as they enter and exit the pluripotent state. However, the field lacks the ability to stably monitor these pluripotency expression dynamics during self-regulation and differentiation in real time. Such reagents would allow the field to properly evaluate current methods for cellular differentiation to increase efficacy, improve our understanding of pluripotency, and elucidate real-time expression dynamics during these developmental processes. Here, we address the gap in tools to elucidate pluripotent transcription factors by providing an efficient and gentle method to create stable monoclonal CRISPR/Cas9 gene-edited hESC lines with OCT4, SOX2, and/or NANOG fluorescently tagged utilizing microraft array technology^46,47^. Our work establishes a robust method for generating live-cell reporters in human pluripotent stem cells lines that will be useful to the human stem cell field. With these reporter tools, we describe real-time heterogenous and morphologically-induced dynamics of these pluripotent transcription factors during multi-lineage differentiations from the developmental perspective of gastrulation and from a single-cell perspective for germ layer differentiations. Our work describes a robust method for generating live-cell reporters in human pluripotent stem cells lines that will be useful to the human stem cell field.

## Results

### Gene editing and clonal selection on microraft arrays

We used CRISPR/Cas9-mediated gene editing to fuse a unique fluorescent protein to each of the three core pluripotency transcription factors in H9 human embryonic stem cells (hESCs)^48–50^ (**Figure 1A-C**; **Experimental procedures**). The three transgenes displayed varying degrees of integration efficiencies, which improved using ribonucleoprotein (RNP)-mediated Cas9 targeting as opposed to plasmid-based targeting (**Figure S1**). Microraft arrays^46,47^, an established technology utilizing magnetic cell culture elements for single-cell sorting applications, were used after gene targeting to identify, isolate, and expand individual stem cells into monoclonal colonies (**Figure 1D**). A key feature of the microraft arrays is that the adhered and spatially separated cells can exchange media between adjacent cells, creating a colony-like environment (**Experimental Procedures**). Immediately after seeding electroporated cells, we performed time-lapse imaging of the full microraft array to identify microrafts that captured a single, fluorescently tagged stem cell. After one week of single cell expansion, microrafts that originally contained a single clone were ejected from the array using a microneedle and transferred to a 96-well plate using a magnetic wand. Clonal colonies were allowed to expand and eventually migrate out from the microraft onto the well plate surface (**Figure 1E**).

**Figure 1.**
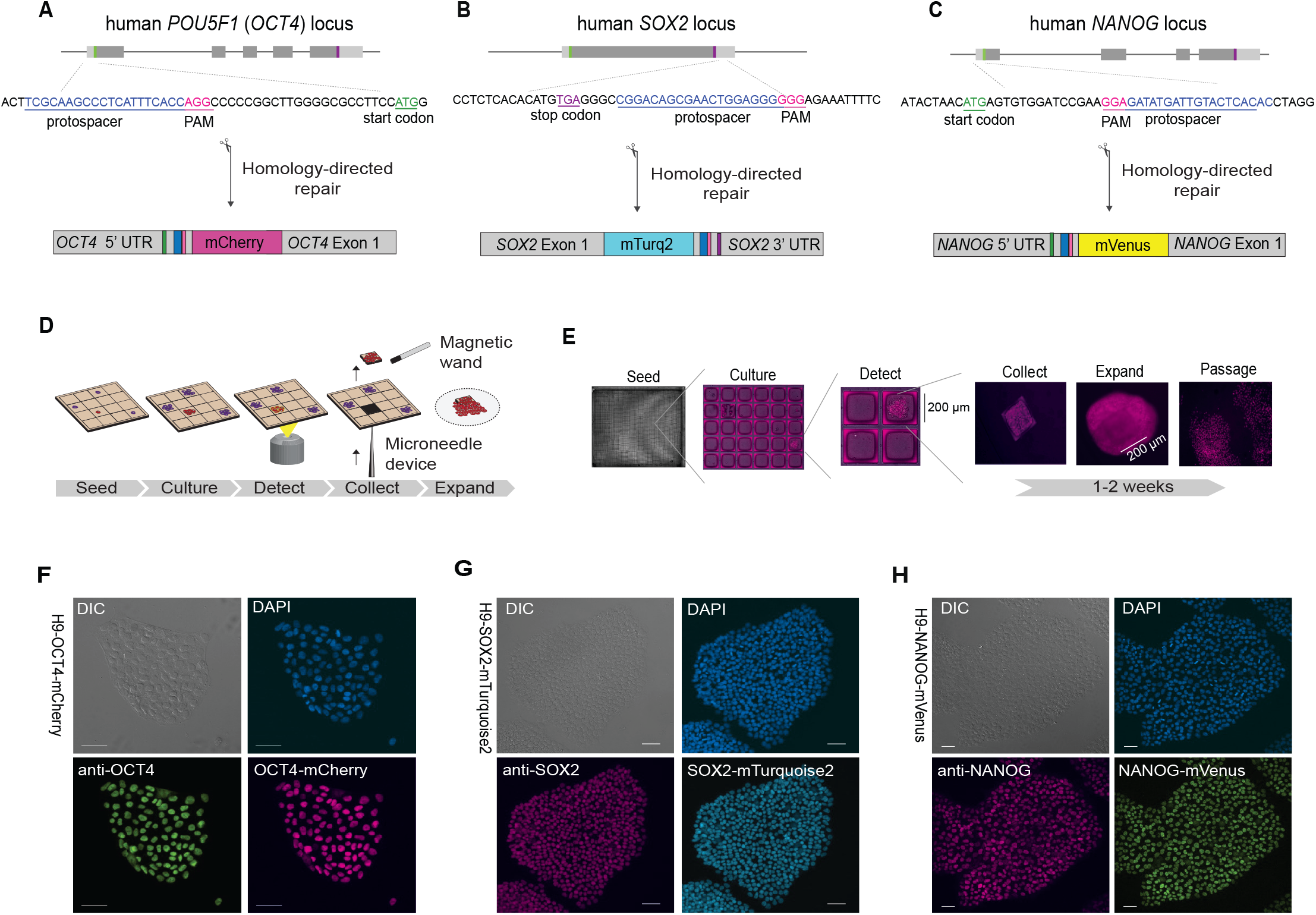
Generation of pluripotent stem cell reporter lines using microraft arrays. **A-C.** Schematic of integration design of the fluorescent protein coding sequence into the endogenous loci of core pluripotency transcription factors in H9 hESCs using CRISPR-mediated homology-directed repair (A, OCT4-mCherry, B, SOX2-mTurquoise, C, NANOG-mVenus). **D.** Schematic of microraft array-enabled cloning procedure. After electroporation of transgenes, hPSCs were seeding onto microraft arrays and imaged by fluorescence time-lapse microscopy to capture early clonal events. After one week, microrafts containing colonies originating from clonal events were ejected using a microneedle and collected using a magnetic wand. The released microraft and its attached cells were deposited onto standard well plates for additional expansion. **E.** Representative images showing identification and expansion of a clonal H9 OCT4-mCherry cell line. **F-G.** Correspondence between expression of endogenously tagged protein and antibody-based immunofluorescence imaging (F, OCT4-mCherry, G, NANOG-mVenus, H, SOX2-mTurquoise).

After culturing multiple clones isolated from individual microrafts, we selected a single stable clone for each of the three core pluripotency transcription factors (OCT4-mCherry, SOX2-mTurquoise2, NANOG-mVenus). These three clones were then subjected to rigorous validation. We confirmed that each transgene showed correct genomic integration, the expected protein fusion size, and a tight correlation with endogenous protein expression (**Figure S2**, **Table S1**, **Figure S3**, and **Figure 1F-H**). Next, we performed whole genome sequencing and karyotyping of the parental H9 cell line and the three reporter cell lines to determine the extent to which gene editing and clonal selection introduced any off-target mutations or chromosomal abnormalities. We observed no significant off-target effects in either gene regions or in potential CRISPR off-target sites and no significant chromosomal changes in the karyotyping (**Table S2, Figure S4**).

Finally, to determine whether gene targeting and fluorophore integration affected the pluripotency or differentiation potential of the hESCs, the reporter cell lines were differentiated into the three primary germ layers: neuroectoderm, mesoderm, and definitive endoderm. All reporter cell lines were capable of differentiation into the three germ layers as determined by transcription of key developmental markers^51^ (**Figure S5**). Thus, the OCT4-mCherry, SOX2-mTurquoise2, and NANOG-mVenus cell lines showed correct genomic integration, accurate reporter protein expression, and proper functional response to developmental cues. We also generated a dual reporter cell line containing both SOX2-mTurquoise2 and NANOG-mVenus for simultaneous monitoring of two pluripotency transcription factors (**Figure S3**, **Table S2**).

### Real time self-organization of hESCs

Having previously characterized the single-cell dynamics of OCT4 in a prior study^52^, we selected the dual reporter cell line containing both SOX2-mTurquoise2 and NANOG-mVenus for further analysis. We first examined the dynamics of SOX2 and NANOG under standard pluripotent growth conditions by live time-lapse confocal microscopy (**Experimental Procedures, Movie S1-2**). Initially, SOX2 and NANOG showed a uniform expression pattern among individual cells within the colonies that formed during the first two days of passaging. However, once the colonies reached a diameter of ~500 μm, the expression patterns changed both in intensity and location within the colonies. Initially, at 300 μm diameter, NANOG decreased in cells located within the center of the colony. After approximately 7 h, the colony grew to ~500 μm in diameter and SOX2 expression increased in cells located within the center of the colony, while NANOG increased in the cells located along the periphery (**Figure 2A, Movie S2**). We did not observe significant migration of cells during colony growth, consistent with previous literature denoting that unperturbed pluripotent stem cells undergoing gastrulation do not have migratory patterning^53^. Other than the lack of migration, the specific expression patterning of SOX2 and NANOG in a localized gradient manner is consistent with previous studies of pluripotent colonies undergoing gastrulation and axial organization from induced morphogen signaling triggered by bone morphogenetic protein 4 (BMP4) addition^54^. This patterning in unperturbed cells resembles gastrulation, a developmental process in which the induction of the primitive streak and germ layers in combination with anterior-posterior axis formation exhibit a spatially specific expression pattern where neuroectoderm is defined by high SOX2 in the center, and definitive endoderm by NANOG in the outer ring of the colonies.

**Figure 2.**
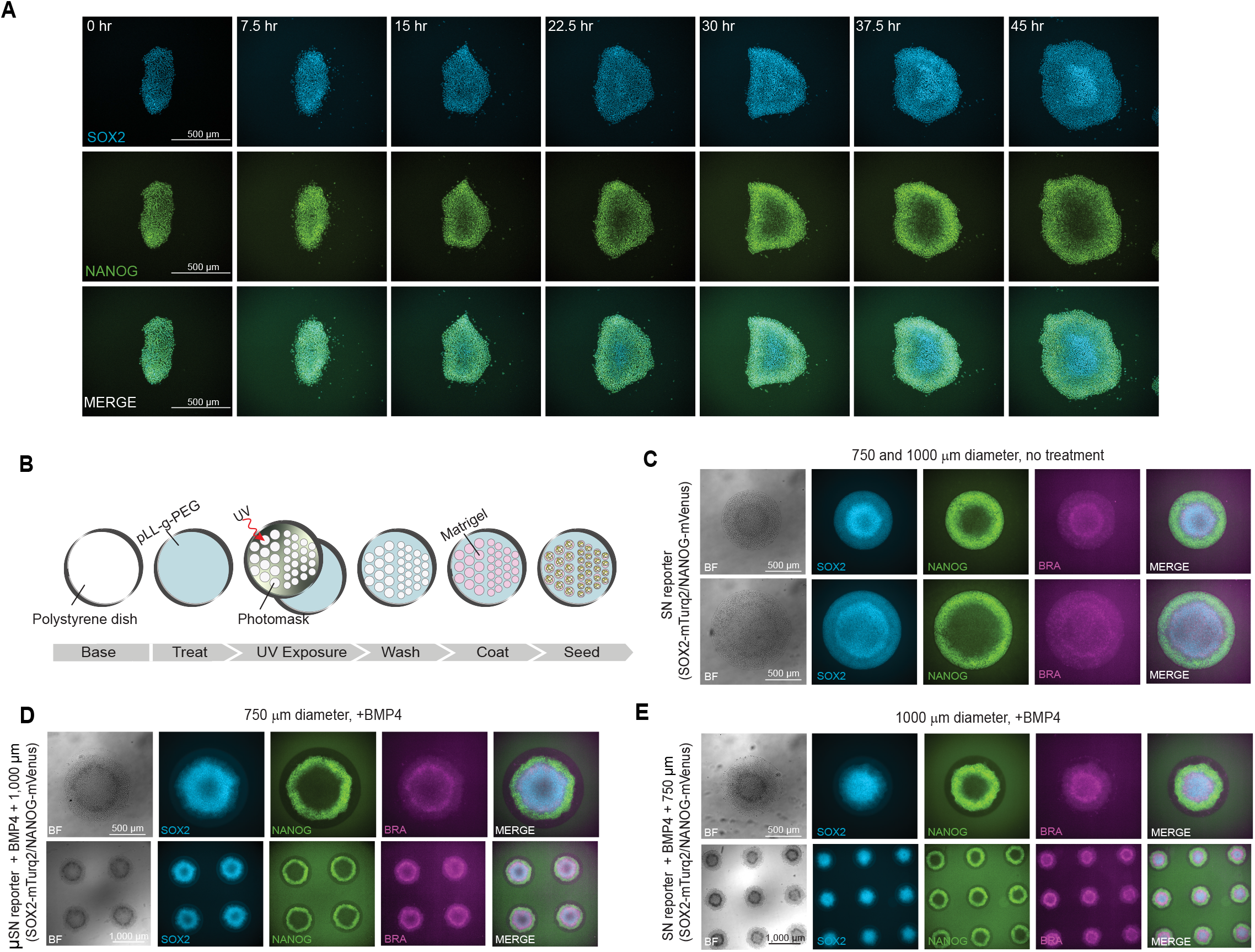
Gastruloid formation and spatiotemporal-specific SOX2 and NANOG expression in live hESCs. **A.** Confocal time-lapse microscopy of simultaneous SOX2-mTurquoise2 and NANOG-mVenus expression in hESC colonues under untreated, pluripotent conditions, showing gastrulation-like expression patterning of SOX2 and NANOG over the course of 45 h. **B.** Schematic of micropattern method to create uniform circular patterns in 750 μm and 1,000 μm diameters. **C.** Dual SOX2-NANOG reporter hESCs 48-hours after seeding without BMP4 treatment on 750 μm (top row) and 1,000 μm diameter (bottom row) micropatterned culture plates with immunoflourescent Brachyury (BRA) antibody staining. **D.** Dual SOX2-NANOG reporter hESCs 48-hours after seeding with BMP4 treatment on 750 μm diameter micropatterned culture plates (top row = 10x magnification; bottom row = 4x magnification) **E.** Dual SOX2-NANOG reporter hESCs 48-hours after seeding with BMP4 treatment on 1000 μm diameter micropatterned culture plates (top row = 10× magnification; bottom row = 4× magnification). C-E. Imaging channels from left to right: brightfield, SOX2-mTurquoise2, NANOG-mVenus, BRA, multi-channel overlay.

To further model gastrulation in live pluripotent cells, we used micropatterning to restrict reporter colonies to a consistent size and shape (**Figure 2B, Movie S3**). When treated with BMP4, these colonies are further induced to form gastruloids^53–55^. Gastruloids have been used successfully to study both cell fate patterning and to identify potential genetic sources of human disease^56–58^. In gastruloids, the dual reporter hESCs showed a similar lack of significant migration while maintaining the same pluripotent transcription factor patterning and expression dynamics between the unperturbed colony growth and micropattern-regulated growth (**Figure 2C**). Treatment with BMP4 was consistent with untreated conditions in patterning but produced a more robust signaling gradient with primitive streak cells expressing higher Brachyury (BRA) expression in an inner ring pattern between the center, delimited by high SOX2, and the periphery, defined by NANOG expressing cells (**Figure 2D-E**). These BMP4-induced gastruloids also showed additional cells present along the edge of the gastruloid that did not express SOX2, NANOG, or BRA, and were not found in the untreated conditions. Previous research utilizing gastruloid models have identified this cell population as trophectoderm-like CDX2+ cells^54^. The formation of trophectoderm-like cells and stronger BRA-positive primitive streak cells suggested that BMP4 addition causes a more rapid induction of gastrulation as well as additional cell fate decisions not present during human embryonic gastrula development.

This well-defined patterning, combined with the observation of limited cell migration, agrees with previous studies showing that changes in cell fate patterning result from signaling between neighboring cells rather than migration of cells with preexisting expression patterns. We directly observed this phenomenon by monitoring two merging colonies expressing the dual reporter (**Movie S1**). The colonies initially grew separately and underwent a gastrulation-like patterning observed in **Figure 2A**. However, within 4 h of the initial contact and fusion into a single colony, NANOG expression along the separate colony edges rapidly changed to encompass the edges of the combined colony, while cells at the newly formed colony junction increased in SOX2 expression. Importantly, the fluorescence signal of both NANOG and SOX2 dynamically travel throughout the fused colony without the individual cells migrating or changing location, affirming that axial organization during gastrulation is caused partially by cell-to-cell communication through edge-sensing mechanisms. This spatially-restricted expression of NANOG and SOX2 in cells suggests that edge-sensing triggers a wave of signaling gradient through otherwise static cells, a phenomenon easily captured through the use of a dual live-cell reporter and validated in non-edited pluripotent stem cell lines (**Figure S6**).

In summary, we demonstrate that our live-reporter system can monitor hESC self-organization in real time; that robust gastruloid patterning may be induced with BMP4 on micropatterned surfaces but chemical induction is not required for gastruloid formation; and that time-lapse imaging reveals dynamic changes in pluripotent transcription factor expression that emerges from changes in colony morphology rather than cell migration.

### Spatiotemporal patterns of SOX2 and NANOG expression during differentiation into endoderm, mesoderm, and ectoderm

In the gastruloid model, pluripotent cells simultaneously differentiated into multiple germ layers in a self-organizing colony. To further elucidate the trilineage fate decision, we next sought to study the dynamics of SOX2 and NANOG expression in each germ layer separately. To do this, we performed directed differentiation of the pluripotent dual reporter cells into neuroectoderm, mesoderm, or definitive endoderm. We used well established and validated protocols for neural progenitor cell (NPC), endothelial cell (EC), or lung progenitor cell (LPC) differentiation (**Figure 3**, **Figure S6**, **Experimental Procedures**). We then used time-lapse fluorescence microscopy to monitor differentiation down each germ layer for 60 h (**Movie S4-6**). Automated segmentation of nuclei was used to quantify nuclear SOX2 and NANOG expression levels every 5 minutes.

**Figure 3.**
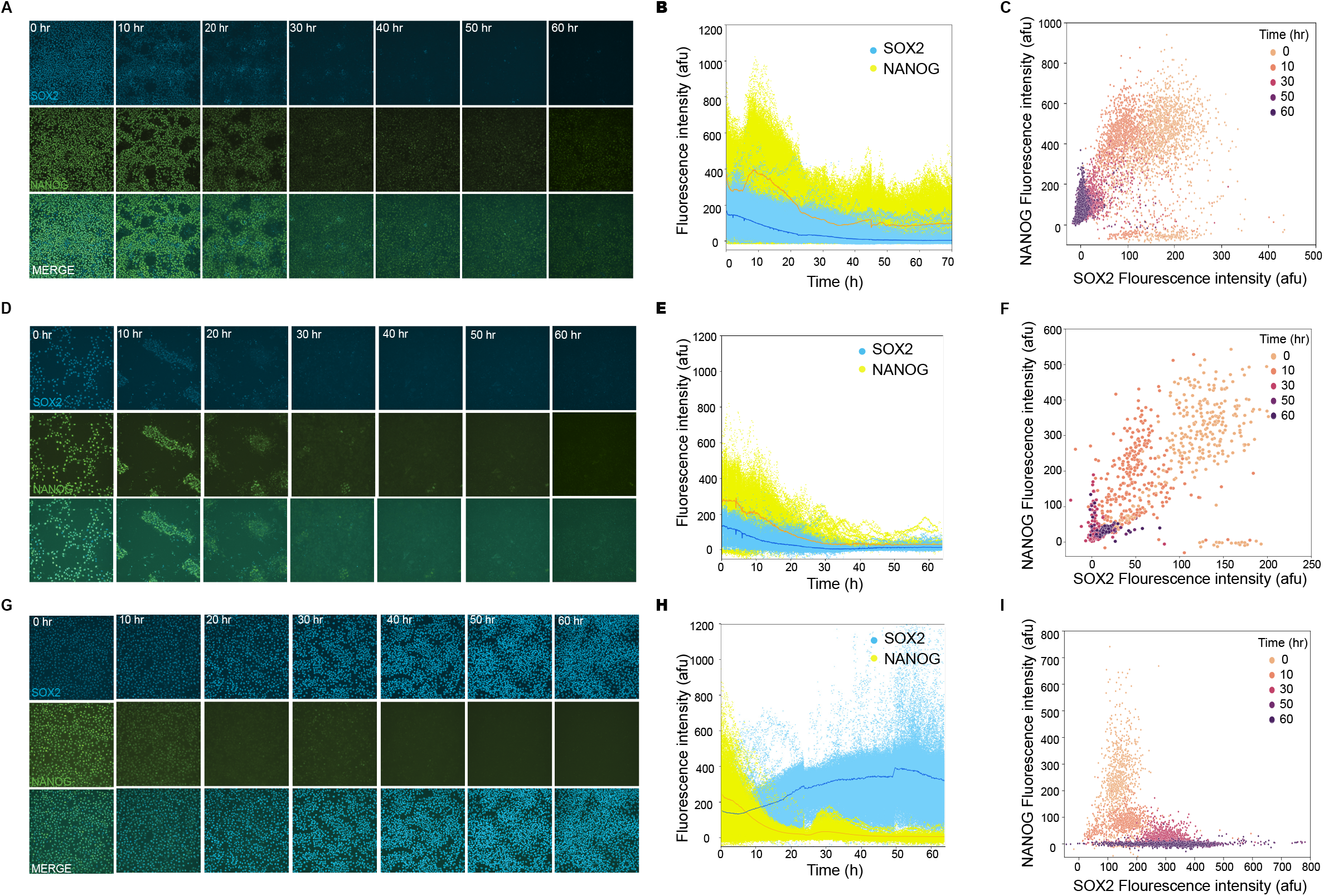
Single-cell dynamics of SOX2 and NANOG protein expression during differentiation into endoderm, mesoderm, and ectoderm. **A.** Film strip of dual reporter (SOX2-mTurquoise2, NANOG-mVenus) expression during lung progenitor cell (endoderm) differentiation. **B.** Single-cell image cytometry of SOX2 and NANOG nuclear intensity during endoderm differentiation. **C.** Single-cell image cytometry of SOX2 and NANOG nuclear intensity at varying time points during endoderm differentiation. **D.** Film strip of dual reporter expression during endothelial (mesoderm) differentiation. **E.** Single-cell image cytometry of SOX2 and NANOG nuclear intensity during mesoderm differentiation. **F.** Single-cell image cytometry of SOX2 and NANOG nuclear intensity at varying time points during mesoderm differentiation. **G.** Film strip of dual reporter expression during neural progenitor cell (ectoderm) differentiation. **H.** Single-cell image cytometry of SOX2 and NANOG nuclear intensity during ectoderm differentiation. **I.** Single-cell image cytometry of SOX2 and NANOG nuclear intensity at varying time points during ectoderm differentiation. >50,000 nuclear images per condition. afu, arbitrary fluorescence units.

Differentiation down the endoderm and mesoderm lineages showed similar temporal dynamics of SOX2 expression (**Figure 3A-F, Movie S4-5**); SOX2 expression decreased to approximately half-maximal levels within the first 12 h of induction. Roughly half of the NANOG signal was lost within ~20 h of differentiation induction in both lineages as well. NANOG showed a similar downward trend except in definitive endoderm, NANOG had fluctuations in expression. Initially, NANOG steeply increased at 5 h, followed by a minor increase at 40 h and a slight increase by 60 h while SOX2 was completely silenced after 30 h in both lineages, NANOG was silenced shortly after for mesoderm but still lowly expressed in a subset of cells in definitive endoderm. Although these changes in transcription factor expression were consistent among most cells, subsets of cells that maintained higher NANOG and SOX2 expression were also present at later time points (**Figure 3C** and **3F**).

In contrast, we observed a different spatiotemporal patterning during differentiation into ectoderm (**Figure 3G-I, Movie S6)**. SOX2 expression increased steadily beginning 5 h after induction and persisted over the course of the differentiation. Similar to the mesoderm differentiation, NANOG expression was silenced within 30 hours. However, we observed a transient reactivation of NANOG in a subset of cells after ~24 h (**Figure 3H** and **3I, Figure 4A**). This reactivation lasted ~6 hours, after which the fluorescent signal was lost. During this time, SOX2 expression continued to rise. Overall, these results indicate hESCs exhibit unique heterogenous SOX2 and NANOG expression dynamics under endoderm, mesoderm, and ectoderm lineage including transient reactivated expression of NANOG under ectoderm differentiation.

**Figure 4:**
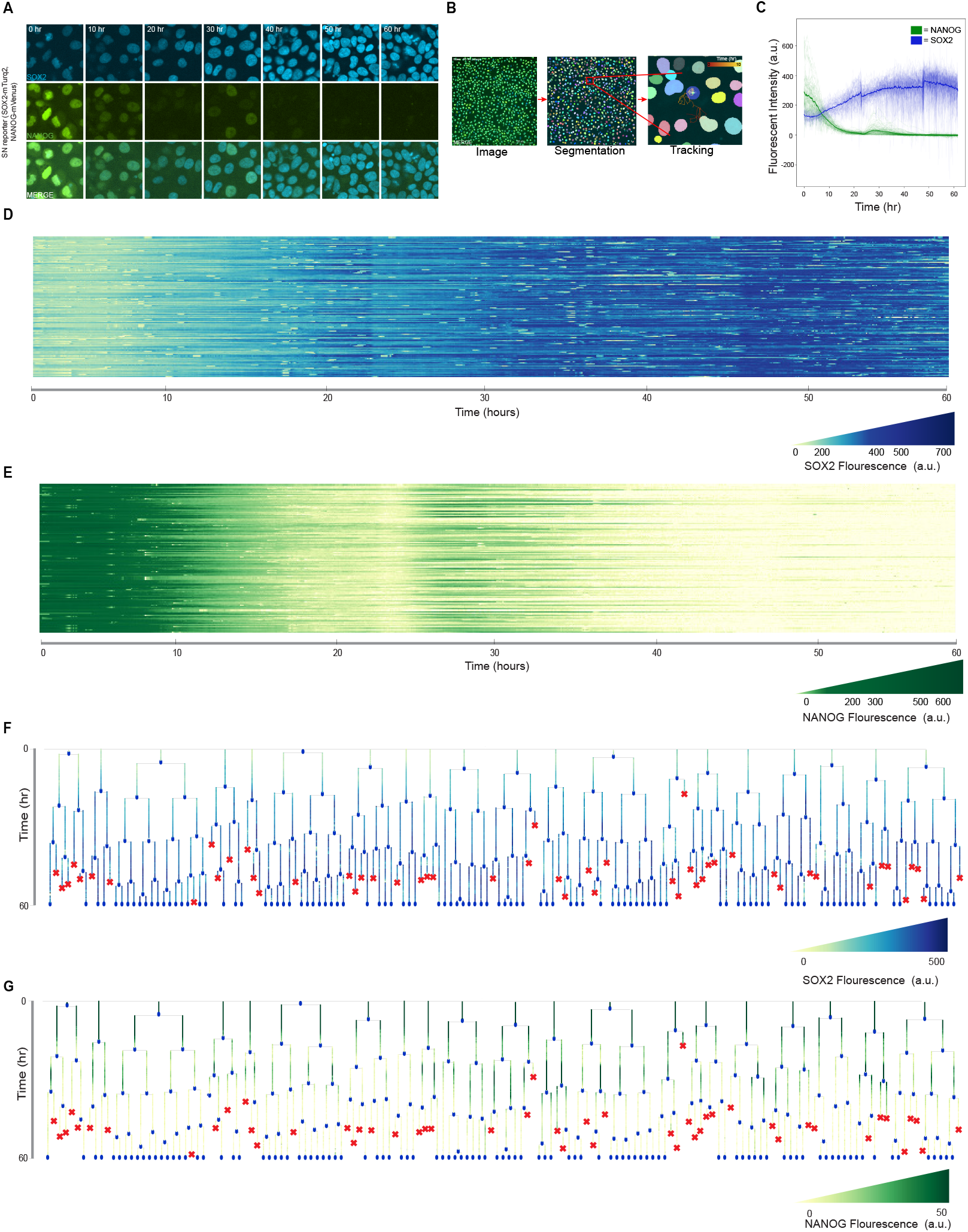
Single-cell dynamics of SOX2 and NANOG expression under differentiation into neural progenitor cells. **A.** Filmstrip of SOX2-mTurquoise2 and NANOG-mVenus expression in live hESCs during differentiation into neural progenitor cells. **B.** Schematic of cell segmentation and tracking algorithms used to quantify mean nuclear fluorescence of SOX2 and NANOG in single cells. **C.** Time series of flourescence intensities of SOX2 (blue) and NANOG (green) of each cell tracked during live imaging. **D.** Lineage tree of full families tracked during live imaging of NPC differentiation with NANOG flourescent intensity range (a.u.). **E.** Lineage tree of full families tracked during live imaging of NPC differentiation with SOX2 flourescent intensity range (a.u.). **F.** Heatmap of non-apoptotic cell lineages during live imaging of NPC differentiation with NANOG flourescent intensity range (a.u.). **G.** Heatmap of non-apoptotic cell lineages during live imaging of NPC differentiation with SOX2 flourescent intensity range (a.u.).

### Single-cell dynamics under differentiation into neural progenitor cells

The previous analysis examined single-cell heterogeneity at regular time points throughout differentiation but did not follow the same individual cells longitudinally over time to assess their single-cell dynamics. Thus, we tracked individual cells over time by linking together expression levels from one frame to the next to elucidate single-cell features. For this analysis, we focused specifically on the ectodermal lineage because of the consistent expression of SOX2 throughout differentiation, which allowed us to clearly follow individual nuclei over the course of the entire experiment (**Figure 4A**). We combined the same segmentation algorithm used to quantify the mean nuclear intensity with a cellular tracking algorithm to understand the differences in expression between single cells and their family lineages (**Figure 4B**, **Experimental Procedures**). Single-cell dynamics for 505 cells showed a general trend of simultaneously increasing SOX2 and decreasing NANOG levels (**Figure 4C**). We again detected NANOG reactivation in a subset of cells ~24-30 h after induction as observed in tracked population-level dynamic analysis (**Figure 4E-F**).

To follow SOX2 and NANOG dynamics over multiple cell generations, we randomly selected 20 cells present at the beginning of the differentiation and tracked cell division events for up to five generations, resulting in a total of 308 cells tracked in the completed family lineages. The fluorescence intensity profiles for the 20 ancestral cell families for NANOG and SOX2 are shown in a full multi-family lineage tree along with apoptotic events, which occurred in 16% (50/308) of cells (**Figure 4F-G**).

### Dynamics of morphological changes during neural progenitor cell differentiation

Throughout the neuroectoderm lineage differentiation, the population underwent morphological changes that indicated cell fate decisions towards neural progenitor cells. Previous work along with our live imaging has shown that neural progenitor cells adopt an elongated morphology as they form near neural rosettes (**Figure 5A**). In response, we tested whether the live imaging system could detect these fate-related morphological changes over time. We fit each nucleus, in each frame of the movie, to an ellipse to quantify the cell’s axial ratio, a measure of how elongated it is (**Figure 5B**). In addition, we also quantified the cell’s orientation as determined by the angle of the major axis with respect to the absolute horizontal direction of the field of view (**Figure 5C**).

**Figure 5:**
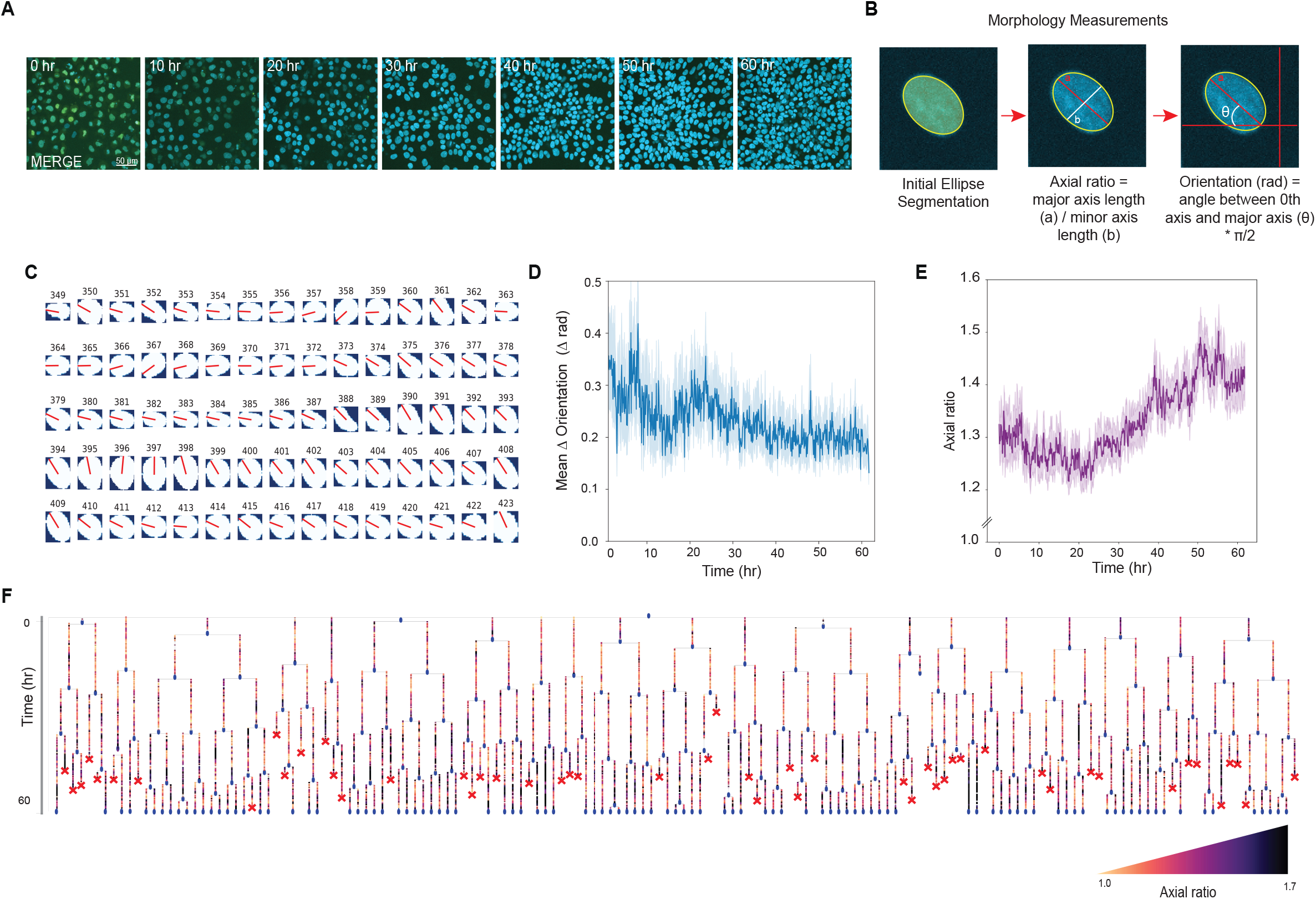
Morphological changes of axial ratio and orientation during NPC Differentiation. **A.** Filmstrip of rosette formation with SOX2-mTurquoise2 and NANOG-mVenus expression during differentiation into neural progenitor cells. **B.** Schematic of how morphological measurements of axial ratio and orientation were created from segmentation algorithm. **C.** Example orientation traces of a single cell’s lifetime over the course of 74 frames (1 frame = 5 minutes) from Frame 349 - 423 (29 h - 35.3 h). **D.** Graph of mean orientation differentiatial over time with 95% confidence interval shown during live imaging of NPC differentiation. **E.** Graph of axial ratio over time with 95% confidence interval shown during live imaging of NPC differentiation. **F.** Lineage tree of full families tracked during live imaging of NPC differentiation with axial ratio range [yellow = 1.0 axial ratio, purple =1.7 axial ratio].

We first quantified the first derivative of the absolute cell orientation to assess how stable the orientation of the cells was over the course of differentiation. We found that the orientation differential decreased throughout the differentiation, reaching a minimum around 45 h after induction, approximately the same time that SOX2 expression plateaued (**Figure 5D**, **Figure 4C**). This shows that the orientation of the cells stabilizes over the course of ectoderm differentiation as the cells determine their positioning to form neural rosettes. Next, we quantified nuclear axial ratio over time in individual hESCs. After a slight decrease in axial ratio during the first 24 h, we observed a steep increase in ovular shape throughout the remainder of the differentiation as cells acquired a more elliptical morphology (**Figure 5E**). A full multi-lineage tree of the axial ratio of the initial 20 ancestral cells throughout the differentiation showed rapid and periodic changes in the elliptical morphology of individual cells, with a gradual trend toward more elongated cells (**Figure 5F**). Thus, the live-cell reporter systems were capable of capturing not only the expression dynamics of pluripotency transcription factors but also morphological features of individual cells, which has been linked to cell fate decisions.

## Discussion

Here, we used microraft arrays to generate clonal CRISPR/Cas9-edited reporter cell lines for real-time monitoring of the pluripotent transcription factors OCT4, SOX2, and NANOG, in hESCs. We are the initial implementors in using microraft array technology for developing reporters for pluripotency transcription factors in human stem cells^46,47^. In contrast to methods based on flow sorting or single-cell well plate seeding, our approach provides a gentle process for clonal selection. The pluripotent cells are not subjected to significant shear stresses, and the individual microrafts are not fully isolated from each other, allowing shared medium between cells for extracellular communication to promote regeneration and survivability^59–62^. We demonstrate that this method can be used for a wide range of CRISPR/Cas9 gene editing, including in induced pluripotent stem cells, and in the generation of a SOX2 biallelic HESC line with different fluorescent tags on opposite alleles of the *SOX2* gene (**Figure S7**). This latter reagent could prove useful in studying genes that undergo allelic switching or are only paternally or maternally expressed^63–65^.

We focused primarily on SOX2 and NANOG protein expression in individual hESCs, monitoring single-cell expression at high temporal and spatial resolution. We observed changes in SOX2 and NANOG expression near colony edges. The edge-sensing morphological influence on differentiation—including during anteroposterior axis formation and control of developmental size—has been reported during embryogenesis in multiple developing organisms^18,66^. Our reporter system highlights the dynamic spatial awareness of hESCs, which were observed to initiate gastrulation-like differentiation without any stimuli or spatial restriction. It has been known and accepted that growth of pluripotent stem cells in pluripotent media generates morphologically identifiable neural rosettes if the pluripotent population becomes over-confluent^67–70^. Our results show that previously termed spontaneous differentiation may instead be caused by the initiation of gastrulation in colonies due to edge-sensing, suggesting that neural rosettes may arise from the center neuroectoderm cells that form within the gastrulating colony. This edge-sensing driver could additionally explain why differentiations have high sensitivities to seeding densities and colony size before the differentiation stimulus is given^71,72^.

The dual reporter also revealed dynamic self-organization of hESC into gastruloids on micropatterns surfaces. By allowing the cells to undergo unperturbed gastrulation, we observed sequential pluripotent transcription factor expression dynamics during this developmental process, with NANOG detected to have an expression and localization change initially before SOX2. Additionally, BMP4-induced micropatterns caused differentiation of extra-embryonic-like cell types not observed in unperturbed cells^54,73,74^. With the ability to observe spatiotemporal changes in germ layer expression, our system may provide an alternative approach to studying gastrulation on micropatterned surfaces without chemical stimuli.

During differentiation toward specific germ lineages—endoderm, mesoderm, and ectoderm—we found that the timing, duration, and intensity of SOX2 and NANOG expression varied between differentiations and within hESC subpopulations in each differentiation. During definitive endoderm, for example, NANOG had multiple fluctuations during its overall downward trend in subpopulations of cells. In neuroectoderm, NANOG expression was lost but showed a brief reactivation in a subset of cells. Meanwhile, SOX2 expression was rapidly lost during mesoderm; lost gradually in subpopulations in definitive endoderm differentiation; and gradually increased during neuroectoderm differentiation. These alternative SOX2 dynamics have been previously reported^10,12,15,34^ but not at the single-cell level nor paired with simultaneous NANOG expression.

One potential explanation for the fluctuations in NANOG expression during definitive endoderm differentiation is the adoption of an intermediate cell fate of mesendoderm^15,17,19^. NANOG is initially expressed in mesendoderm cells and drives the eventual bifurcation into definitive endoderm. NANOG and co-activator GATA6 promote definitive endoderm fates and remodel chromatin to repress pluripotent plasticity, but NANOG increases only briefly in mesoderm before repression as to diverge from pluripotency, but not undergo full endoderm differentiation^10,75^. Further, NANOG is a main driver in endoderm fate decisions and binds with various fate-specific transcription factors, such as GATA6, throughout the differentiation to drive loss of pluripotency, suppress ectoderm and mesoderm fates, and promote endoderm formation from mesendoderm intermediates^23,76,77^. These essential roles could translate into initial bursts of NANOG expression being a necessity for definitive endoderm cell fate specification as it could potentially cause prolonged expression of NANOG to allow for NANOG to bind to SMAD2/3 and recruit EOMES during a mesendoderm intermediate transition.

For ectoderm, the brief reactivation of NANOG observed during ectoderm differentiation coincided with changes in morphology toward the cell fate decision of neuroectoderm. It is well-documented that morphological changes have a direct effect on certain cell fate decisions, and that the morphology of a given cell or within a tissue can cause variable somatic cell type fate decisions^78–80^. Our data showed NANOG recapitulation occurred before major morphological and orientation changes such as the formation of early neural rosettes^81–83^, which derives interest of whether NANOG is involved with morphological cues or has an indirect role. There is evidence to support that nuclear β-catenin is associated with NANOG increase^84,85^; β -catenin’s location throughout the nuclear envelope is required for neural differentiation as well as morphological changes such as epithelial-to-mesenchymal (EMT) transitions^86–88^. NANOG could potential play a role in accumulating β -catenin to cause neuroectoderm fates and eventual NPC specification and morphology. Further work is needed to determine whether NANOG reactivation is a predictor of hESC morphology during differentiation of neural progenitor cells.

Heterogeneity in pluripotent transcription factors, particularly NANOG, has been previously observed along multiple cell fate decisions^89–91^. This heterogeneity may provide a stochastic advantage during differentiations—especially between naïve and primed stem cells—by providing more flexibility to adopt different cell fates^92–94^. Since NANOG is required to maintain pluripotency but cannot necessarily induce it^95,96^, it is a potentially more sensitive and variable pluripotent TF compared to SOX2 or OCT4 which could add another factor in its multi-lineage heterogeneity.

In summary, we describe a novel and efficient method for generating reporter cell lines in human pluripotent stem cells and provide insights into the single-cell dynamics of SOX2 and NANOG during differentiation into the three primary germ layers. Our work shows that the dynamics of these transcription factors are highly variable and rapidly changing as cell fates are acquired. It is possible that this heterogeneity could be exploited to improve differentiation efficiency or to predict a successful cell fate decision.

## Experimental procedures

### Resource Availability

Corresponding authors JEP and ASB have listed materials, code, and data available from the lead contact without restriction.

### Human embryonic stem cell culture

The human embryonic stem (hES) cell line H9 (WA) was obtained from WiCell. H9 and stable reporter OCT4, SOX2 and NANOG were maintained on Matrigel (Corning) in StemFlex medium (Thermo Fisher Scientific) and routinely passaged every 3 to 4 days with 0.5 mM EDTA/PBS dissociation solution at 1:6 ratio. After passage, cells were cultured in the presence of 10 μM Y-27632 (ROCK inhibitor, Peprotech) for 24 hours in a humidified incubator at 37°C and 5% CO_2_.

### CRISPR/Cas9 genome editing

We used the Cas9/gRNA ribonucleoprotein (RNP) method and the Neon™ Transfection system (Thermo Fisher Scientific) to edit the H9 cell line. Recombinant TrueCut Cas9 V2 (Thermo Fisher Scientific) was diluted in resuspension buffer R, mixed with specific, modified sgRNAs (Synthego), and incubated for 15 minutes at room temperature. The following modified sgRNAs were utilized in these experiments: OCT4: 5’-gUgaaaUgagggcUUgcga-3’, NANOG: 5’-cacacUcaUgUUagUaUag-3’ and SOX2: 5’-gacagcgaacUggaggg-3’. A total of 3×10^5^ cells, 1×10^5^ cells in each 10 ul Neon™ Transfection tip per electroporation reaction repeated three times, were electroporated with 900 μg sgRNA, 3,000 ng linearized donor plasmid, and 3,000 ng Cas9 enzyme using the Neon program 17 (850v/30ms width/2 pulses). After 48 to 72 hours, integration of the donor plasmid was confirmed by fluorescent microscopy. After 96-hours, cells were dissociated into single cells and 6000 cells were seeded onto the microraft arrays.

### Microraft Array Generation

Microraft arrays were fabricated according to previously reported protocols^97^. Each array was made to be 21 × 21 mm and comprised a total of 12,000 magnetic microrafts, with each microraft having a 200 × 200 × 100 μm geometry with 30 μm of space between each microraft. Select microraft locations across the array (every 10^th^ row and column) possessed a numerical code, enabling microraft identification and tracking. Microraft arrays having this geometry are also sold commercially as CytoSort™ arrays and can be purchased from Cell Microsystems (Durham, NC).

### Clone Identification and Collection

To augment the microraft arrays fabricated in-house, a CellRaft™ system was obtained from Cell Microsystems and used to release magnetic microrafts containing adhered target colonies from the array. Key components of the system include a release device, control box, and magnetic wand. The release device fits reversibly onto the objective of an inverted microscope and has a microneedle mounted to the top. The control box is used to set the height that the microneedle will need to move to touch the microraft array positioned on the stage, thus causing a target microraft to be dislodged from the array. Microrafts are easily collected with the magnetic wand and deposited into a well plate containing cell culture medium for continued culture of the adhered cells.

To produce monoclonal hPSC colonies, cells were seeded as single cells onto a microraft array and cultured into microcolonies. The entire microraft array was scanned using fluorescence microscopy immediately after cells were seeded and as colonies formed. Image analysis was used to identify target microrafts possessing a single cell and its resulting monoclonal colony stably expressing the fluorescent reporter. Stability was denoted as fluorescence expression in all cells and consistent pluripotent cell morphology with little to no stress morphology of spiked cytoplasm present. Microrafts with target colonies were released from the array, deposited into a well plate, and the adhered colony allowed to migrate off the microraft and onto the well plate surface. Standard protocols were used for continued culture and expansion of the cells.

### Sequencing of cell lines

DNA was extracted and purified using (Zymo gDNA kit) according to the manufacturer’s recommendations. Extracted DNA was quantified using the Qubit DNA Assay Kit and a Qubit 2.0 Flurometer (Life Technologies, CA, USA). Genomic DNA was sheared to 350bp by sonication and 1.0 ng of each sample was used for sequencing library generation (NEBNext DNA Library Prep Kit). Libraries were sequenced on an Illumina NovaSeq. Reads were then filtered for adapter contamination using cutadapt (Martin, 2011) and filtered such that at least 90% of bases of each read had a quality score >20. Duplicated sequences were then capped at a maximum of 5 occurrences, and reads were aligned to the reference genome (hg19) using STAR ((Dobin et al., 2013)) version 2.5.2b retaining only primary alignments.

### Whole Genome Sequencing-Table 1

Allele read depth was analyzed to ensure SNPS had at least 15 reads with at least 5 reads for each called allele. Excluded SNPs on unassembled contigs.

### Variant calling

Variants were identified using the Genome Analysis Toolkit (GATK) v3.8 and the developers’ suggested best practices^98^. Briefly, the BAM files had read groups and names modified by sample, and duplicates were marked using Picard tools (http://broadinstitute.github.io/picard/). The read qualities were then recalibrated, and SNPs were called using BaseRecalibrator and HaplotypeCaller, respectively. The resulting VCF files for each sample were merged into a single GVCF containing SNP calls for all samples using CombineGVCFs and GenotypeGVCFs. Variant calls were recalibrated using VariantRecalibrator. Variants from Hapmap 3.3, 1000 genomes, and dbSNP 138 were used to build the variant recalibration model. The variants in the final VCF were annotated for associated genes and severity/function using snpEff v4.3^99^. Variants were then filtered to retain those denoted as “PASS”, containing a read depth of at least 15 in all samples, a read depth of at least 5 for each called allele in all samples, and then variants identified in all five samples were removed. We also called variants with an independent approach using Strelka v2.9.10^100^. Variants were called for all reporter cell lines relative to WT. The union set of variants were then used to intersect with those identified by GATK to obtain one set of high-confidence variants identified by both methods.

### Characterization of pluripotency and differentiation capabilities

To confirm stemness and differentiation capabilities of the reporter lines we used the qPCR based TaqMan hPSC Scorecard Panel (ThermoFisher Scientific). H9 cells were differentiated into all three germ layers using STEMdiff Trilineage Differentiation Kit (StemCell Technologies), a mono-layer based protocol to directly differentiate hES cells in parallel into the three germ layers. RNA extraction, quantification and reverse transcription for both undifferentiated and differentiated hES cells was performed as described above (or in the *quantitative real-time polymerase chain reaction (qRT-PCR) section).* Taqman Scorecards were processed according to manufacturer guidelines using the QuantStudio 7 Flex Real-Time PCR system. Scorecard results were exported into the TaqMan hPSC Scorecard Panel online data analysis platform that compares the input gene expression profile to a common reference set. The TaqMan PCR assay combines DNA methylation mapping, gene expression profiling, and transcript counting of lineage trace marker genes^101^. After an intuitive analysis, the software automatically reports if the sample consists of cells with stemness or differentiated properties.

### Immunofluorescence staining

Undifferentiated and differentiated cells (including gastruloids) were fixed with 4% paraformaldehyde (ThermoFisher Scientific) for 15 minutes, then permeabilized with 0.03% Triton X-100 and blocked with 5% bovine serum albumin (Sigma-Aldrich) for 2 hours at room temperature. Plates were then incubated in primary antibody solution overnight at 4 °C, followed by an incubation with secondary antibodies for 1-2 hours at room temperature. Primary antibodies used in this study included Anti-FOXA2 antibody [EPR4466] Rab mAb, Anti-SOX17 antibody [EPR20684] mouse pAb, Anti-Brachyury / Bry antibody [EPR18113] Rab mAb, anti-PAX6 antibody (42-6600) Rab pAb, anti-SNAIL antibody (C53895) mouse mAb, anti-SOX1 antibody (MA53244) Rab mAb, anti-SOX2 antibody (Ab97959) Rab pAb, and anti-NANOG antibody (ab109250) Rab mAb (used at 1:200-1:1000). Secondary antibodies used were Alexa Fluor 488, 568, 596, and 647 (used at 1:500; Abcam). Cells were counterstained using 4’,6-diamidino-2-phenylindole (DAPI, 1:5000) in 1x PBS, then washed and imaged or stored at 4 °C in PBS.

### Western Blotting

Cells were lysed with Pierce RIPA buffer for 30 minutes then centrifuged at maximum speed for 20 minutes. Supernatant was collected and stored at −80°C until ready for western blotting. Cell lysate was denatured with BioRad 2X Laemmli buffer then separated by SDS-PAGE using a TGX Any KD gel (BioRad, 456-9054). Transfer to PVDF membrane was performed using Trans-Blot^®^ Turbo™ RTA Mini PVDF Transfer Kit and the TGX mini transfer protocol on the BioRad Trans-Blot^®^ Turbo™ transfer machine. Membranes were blocked for one hour at room temperature in Odyssey blocking buffer TBS then incubated overnight at 4°C with primary antibodies at 1:1000 in blocking buffer. Following washing with TBST, membranes were incubated in Licor IRDye secondary antibodies at 1:14000 in blocking buffer for one hour at room temperature. Membranes were washed three times with TBST followed by a wash with TBS and imaged using a Licor Odyssey CLx scanner at 800 and 700 nm. Anti-SOX2 antibody [EPR3131] Rab mAb. Abcam ab92494. Anti-Oct4 Rab pAb. Abcam ab19857. Anti-Nanog antibody [EPR2027(2)] Rab mAb. Abcam ab109250. Anti-histone mouse mAb. Cell Signaling 3638S.

### PCR

Genomic DNA was extracted from each reporter cell line and subjected to PCR with primer sets flanking the targeted gene. PCR products were separated by electrophoresis on 1.5% agarose gels stained with Green Gene Safe DNA Dye (Southern BioLabs). PCR products were purified using Zymo’s DNA Clean and Concentrator kit and sequenced with their respective primer sets to confirm correct insertion of the fluorescent tag.

### Karyotyping

H9 cells and monoclonal reporters were submitted to a standard G-band karyotype analysis consisting of 20 metaphase spreads. The analysis was carried out by Karyologic Inc.

### Micropatterning

The micropatterned culture plates were created by initially treating 6-well plastic culture dishes (Corning) with 1 ml per well of 10 mg/ml poly-L-Lysine graft poly(ethylene glycol) with FITC labeling (pLL[20]-g[3.5]-PEG(2)/FITC; SuSoS Technologies) diluted 1:100 in room temperature PBS for 1 hour at room temperature. After, the solution was fully aspirated, and 15 μl of room temp. DI water was placed in the middle of the well. A 33-mm diameter quartz 10 μm resolution mask (FrontRange Photomasks) with 750 and 1,000 μm diameter circles was placed in the well chrome-side down and exposed to 254 nm ultraviolet radiation for 15 minutes in the HELIOS-500 UV ozone cleaner (UVFAB). Wells that were treated with pLL-g-PEG/FITC but did not have the quartz mask were covered with tinfoil to preserve solution coating. After UV exposure, room temp. DI water was added to the well and the quartz mask was removed using a pipette tip to lift the mask. The well was washed three times with room temp. DI water, and then coated with two ml of Matrigel (1:100 in DMEM/F12) overnight at 4°C. The next day, the Matrigel was slowly diluted by removing one ml of Matrigel and adding one ml of StemFlex basal media (STEMCELL Technologies) six times to not expose coating to air. Finally, one ml was removed and replaced with 2x StemFlex media. This process created a surface with circular patterns of matrigel surrounded with cell-repellent pLL-g-PEG.

### Gastruloid Formation

Gastruloids were created through a micropatterning process where the culturing dish was treated in a method in which the cells would only adhere to the circular patterns of Matrigel (**Figure 1B**, micropatterning methods section). The SOX2/NANOG reporter cells were treated with ROCK inhibitor (10 μM) for one hour before passaging, and two hours after passaging. The cells were passaged with PBS/0.5 mM EDTA, seeded on a micropatterned 6-well plastic culture plate at 500,000 cells/well, and fed daily with StemFlex. The cells were allowed to fill the micropatterns before treatment; this took approximately 48 hours. After they filled these micropatterns, 100 ng/μl of BMP4 (STEMCELL Technologies) was added daily for 48 hours, then fixed for immunofluorescent staining.

### Live Imaging

All live imaging of untreated growth, ectoderm, mesoderm, and endoderm differentiations were collected with Nikon Ti Eclipse A1-A Confocal microscope operated by NIS Elements software V5.02.00 with an Andor ZYLA 4.2 cMOS camera and a custom stage enclosure (Okolabs) to ensure constant temperature, humidity, and CO2 levels at 37°C and 5% CO2.The exposure, excitation wavelength, binning, bit-depth, and power for each channel were kept consistent for all three live imaging differentiations. For each trilineage differentiation live imaging movie, SOX2-mTurquoise2 / NANOG-mVenus H9 cells were initially dissociated with pre-warmed 37°C Accutase Cell Detachment solution (Innovative Cell Technologies, Inc.) for 3-5 minutes, collected with StemFlex (Thermo Fisher Scientific), spun down at 1,000 × g, and resuspended on 12-well plates with a glass-like polymer bottom (Cellvis) pre-coated with Matrigel at 37°C for two hours (Corning).

### Untreated Growth Live Imaging

After dissociation with SOX2-mTurquoise2 / NANOG-mVenus H9 cells with 0.5 mM EDTA/PBS dissociation solution, cells were seeded in a 6-well plate with a glass-like polymer bottom (Cellvis) at a 1:6 ratio. After passage, cells were cultured in the presence of 10 μM Y-27632 (ROCK inhibitor, Peprotech) for 24 hours in a humidified incubator at 37°C and 5% CO_2_, then imaged 48 hours after seeding every 5 minutes with a media change every 24 hours for 48 hours total.

### Neuroectoderm / Neural Progenitor Cell Differentiation Live Imaging

After dissociation, 1.25 million SOX2-mTurquoise2 / NANOG-mVenus H9 cells were resuspended in 1 ml of STEMdiff™ Neural Induction Medium (NIM, Stem Cell Technologies) with STEMdiff™ SMADi Neural Induction Supplement (SMADi) and 10 μM Y-27632 (ROCK inhibitor, Peprotech) (or StemFlex (Thermo Fisher Scientific) with 10 μM Y-27632 (ROCK inhibitor, Peprotech) for negative control, data not shown) and then seeded per well in a 12-well plate. The plate was put in an incubator at 37°C and 5% CO2 for 30 minutes to ensure cell attachment, then transferred to the Nikon Ti Eclipse confocal microscope with enclosure at 37°C and 5% CO2 and imaged for SOX2 and NANOG expression every 5 minutes. Approximately 22 and 45 minutes after movie started at Frame 273 media was changed to warmed NIM with SMADi, at 47 hours and 45 minutes at Frame 572 media was changed to warmed NIM with SMADi, and at 61 hours and 55 minutes at Frame 741 movie was stopped.

### Definitive Endoderm / Lung Progenitor Differentiation Live Imaging

After dissociation and resuspension, 1 million SOX2-mTurquoise2 / NANOG-mVenus H9 cells were seeded per well in a 12-well pre-coated with Matrigel (Corning) for two hours at 37°C with StemFlex medium and 10 μM Y-27632 (ROCK inhibitor, Peprotech). The plate was put in an incubator at 37°C and 5% CO2 for 30 minutes to ensure cell attachment, then transferred to the Nikon Ti Eclipse confocal microscope with enclosure at 37°C and 5% CO2 and imaged every 5 minutes. After 25 hours of imaging (F299), the media was changed to pre-warmed 37°C Medium DE-1 (STEMdiff™ Endoderm Basal Medium containing Supplement MR and Supplement CJ, Stem Cell Technologies). One well of seeded cells was left untreated as negative control (data not shown). At 46.8 h (F562), the medium was changed to Medium DE-2 (STEMdiff™ Endoderm Basal Medium containing Supplement CJ), and at 70.1 h (F841), the medium was changed to Medium DE-2 again. The plate was continuously imaged every 5 minutes until 95.3 h (1144F).

### Mesoderm / Endothelial Progenitor Cell Differentiation Live Imaging

After dissociation and resuspension, 300,000 SOX2-mTurquoise2 / NANOG-mVenus H9 cells were seeded per well in a 12-well pre-coated with Matrigel (Corning) for two hours at 37°C with StemFlex medium and 10 μM Y-27632 (ROCK inhibitor, Peprotech). The plate was put in an incubator at 37°C and 5% CO2 for 30 minutes to ensure cell attachment, then transferred to the Nikon Ti Eclipse confocal microscope with enclosure at 37°C and 5% CO2 and imaged for SOX2 and NANOG expression every 5 minutes. After approximately 24.1 hours of imaging (F289-F290), the media was changed to pre-warmed 37°C mesoderm / Endothelial “priming” media (8 uM CHIR99021 +25 ng/ml BMP4, 1× Glutamax, 1× N2, 1× B27, Neurobasal, DMEM/F12, Stem Cell Technologies). One well of seeded cells was left untreated as negative control in StemFlex (data not shown). The plate was continuously imaged for SOX2 and NANOG expression every 5 minutes until 86.1 h (Frame 1033).

### Image Analysis

Image analysis was performed using Python (3.7.10). Segmentation of cell nuclei was performed with the ‘cyto’ model from Cellpose (v. 0.6.5)^102^. Characterization of objects was performed using Scikit-image (v.0.18.2)^103^ image processing library. Initial linking of the objects into tracks was performed with BayesianTracker (v. 0.4.1)^104^. Errors in segmentation and tracking were corrected manually using Napari (v. 0.4.10)^105^ graphical interface. A total number of 505 cells were tracked and 20 full lineage families. Full lineage families were selected to track by using a random number generator with all Cell IDs at Time 0 then systematically tracking cells by Cell ID list. The first 20 cells that survived past Frame 50 without apoptosis were expanded to track full lineages. This totaled 290 out of the 505 cells tracked. After full families were tracked, cells from Time 0 to apoptosis or end of movie were tracked with incomplete lineages. Graphs of live imaging data were synchronized for Time 0 to reflect differentiation media treatment time. Orientation quantification included assumptions that rotations larger than π/2 between a single frame is due to axial rotation. To construct single-cell heat maps across the length of the movie, we selected all tracked lineages with at least 80% non-missing values in SOX2 and NANOG expression. This resulted in 145 total lineages, each with multiple generations of cells, tracked from the beginning to end of the movie. All processing and analysis were done in R. Missing values are linearly interpolated using the zoo library and heatmaps are plotted using the superheat library in R. Microscopy images were adjusted for brightness and contrast in NIS Elements software V5.02.00 and were uniformly adjusted for trilineage differentiation.

## Supporting information

Highlights and eToc blurb

## Acknowledgements

We thank Jolene Ranek for helping develop Jupyter notebooks for image analysis.

## Author Contributions

SM designed, optimized, and produced micropatterned substrates for gastruloid analysis, carried out differentiation and live imaging studies, tracked individual cells, assisted in validating the reporter cell lines, prepared all main text figures, and wrote the first draft of the paper. SCW and ASB constructed the reporter cell lines: SCW designed and prepared the gene tagging reagents and coordinated the cell line construction, validation, and whole genome sequencing efforts. ASB carried out electroporation, expansion, and maintenance of monoclonal cell lines, and performed validation and differentiation experiments. KMK designed custom image analysis tools and supported single-cell segmentation tracking. NS developed the microraft array technology for use with human stem cells. MR validated reporter cell lines. YW advised on microraft arrays and micropattern optimization. GKL performed single cell tracking. JS and TP performed variant detection in whole genome sequencing. NLA, ASB, and JEP provided supervision and mentorship for the work. This research was supported by NIH grants R01-GM138834 (JEP), DP2-HD091800 (JEP), and R01-CA224763 (NLA); NSF CAREER Award 1845796 (JEP), and Chan Zuckerberg Initiative DAF grant 2020-225716 (KMK).

