## Supplementary material for "Single-cell dynamics of core pluripotency factors in human pluripotent stem cells": Highlights and eToc blurb

### Highlights:

- Microraft array used to establish novel OCT4, SOX2, NANOG single and dual reporters
- Self-organization of untreated pluripotent cells undergo gastrulation-like patterning
- Trilineage differentiations reveal spatiotemporal pattern for SOX2 and NANOG
- Ectoderm lineage has change in nuclear morphology during cell fate commitment

### eTOC Blurb:

Mihailovic et al. have provided the field with a novel robust method for generative live-cell reporters. They've created live-cell reporters of core pluripotency transcription factors and combined the reporters with time-lapse imaging to study self-organization, formation of two-dimensional gastruloids, and the single-cell dynamics of directed tri-lineage differentiation. This revealed unique self-organization, spatiotemporal patterning, and morphological changes.
