## Supplemental Material for "Single-cell dynamics of core pluripotency factors in human pluripotent stem cells"

† Corresponding Authors

**Contents:**

**Figure S1** **Integration efficiency in ribonucleoprotein (RNP) versus plasmid-mediated gene editing**

**Figure S2** **Genomic integration of fluorescent fusion proteins**

**Table S1** **Oligo sequences for primers used in Figure S2**

**Figure S3** **Comparison of endogenous and reporter protein expression**

**Table S2** **Genomic alterations introduced during development of reporter cell lines.**

**Figure S4** **Karyotyping analysis of clonal reporter cell lines.**

**Figure S5** **Transcription profiles of reporter cell lines upon directed differentiation.**

**Figure S6** **Immunofluorescence validation for proper differentiation and unperturbed self-organization.**

**Figure S7** **SOX2-mVenus/SOX2-mCherry (biallelic) cell line validations.**

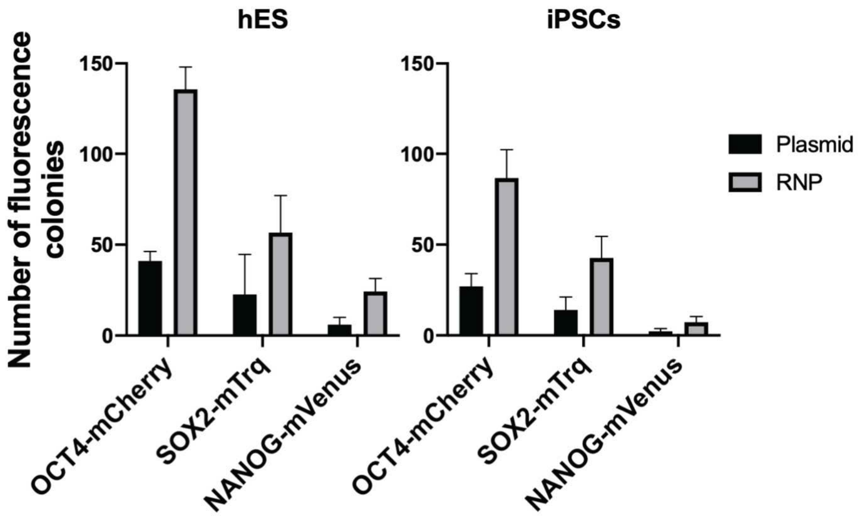


**Figure S1. Integration efficiency in ribonucleoprotein (RNP) versus plasmid-mediated gene editing.** H9 human embryonic stem cells (hESCs) or 15CA induced pluripotent stem cells (iPSCs) were electroporated with plasmid-encoded Cas9 or in translated ribonucleoprotein (RNP) form along with the fluorescent donor cassette (see Materials and Methods). As an approximate measure of efficiency, the number of fluorescent colonies was counted as successful integration events.

A.

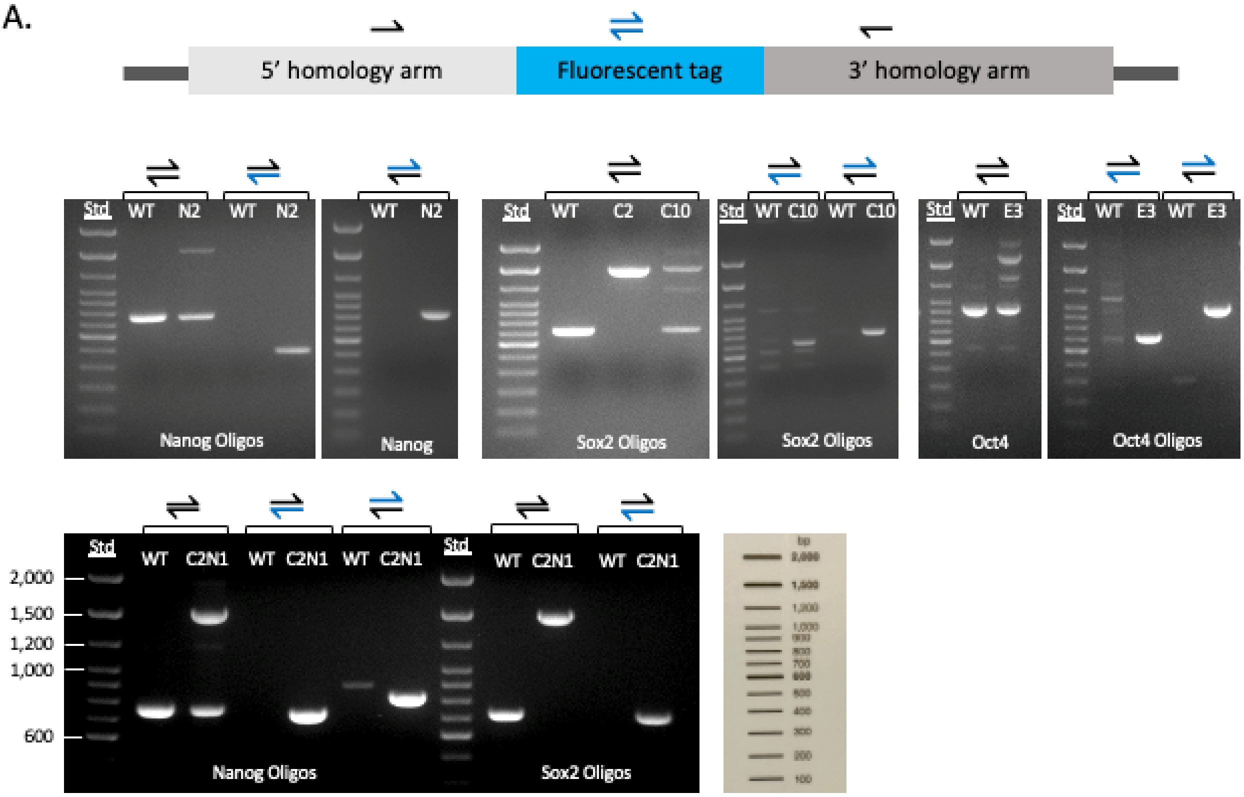
5' homology arm
Fluorescent tag
3' homology arm
Std
WT
N2
WT
N2
WT
N2
WT
C2
C10
WT
C10
WT
C10
WT
E3
WT
E3
WT
E3
Nanog Oligos
Nanog
Sox2 Oligos
Sox2 Oligos
Oct4
Oct4 Oligos
2,000
1,500
1,200
1,000
600
WT
C2N1
WT
C2N1
WT
C2N1
WT
C2N1
WT
C2N1
Nanog Oligos
Sox2 Oligos

B.

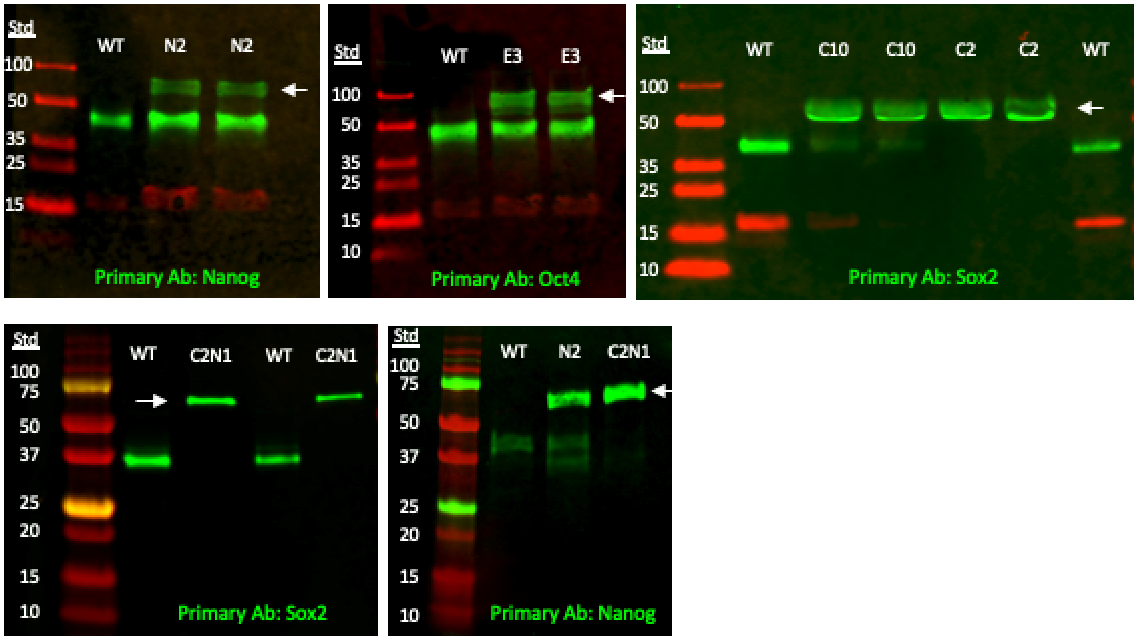
Std
WT
N2
N2
100
50
35
25
15
Primary Ab: Nanog
Std
WT
E3
E3
100
50
35
25
15
10
Primary Ab: Oct4
Std
WT
C10
C10
C2
C2
WT
100
50
35
25
15
10
Primary Ab: Sox2
Std
WT
C2N1
WT
C2N1
100
75
50
37
25
20
15
10
Primary Ab: Sox2
Std
WT
N2
C2N1
100
75
50
37
25
20
15
10
Primary Ab: Nanog

**Figure S2. Genomic integration of fluorescent fusion proteins. A.** Genomic DNA was extracted from each of the indicated reporter cell lines and subjected to PCR with a set of primers flanking the targeted gene. The table below of PCR reactions and correlation with the images above indicates correct insertion of the fluorescent protein gene. **B.** Cell lysate from the indicated reporter cell lines as well as H9-WT cells were run out in duplicate on a gradient SDS-PAGE then transferred and blotted with antibodies for human NANOG, OCT4 or SOX2 protein. Western blot analysis revealed successful integration of the fluorescent protein tag in all the reporter cell lines as indicated by the arrowheads. N2. NANOG-mVenus clone (mono-allelic). C2, SOX2-mTurquoise clone (bi-allelic). C10, SOX2-mTurquoise clone (mono-allelic). C2N1, SOX2-mTurqoise (bi-allelic)/NANOG-mVenus (mono-allelic) dual reporter clone. E3, OCT4-mCherry clone (mono-allelic).

| Oligo Name | Oligo Sequence (5’->3’) | WT Product Size (bp) | FP Insertion Product Size (bp) |
|---|---|---|---|
| Nanog Blk Fwd[1] | CAGGTTCTGTTGCTCGGTTT | 755 [1/2] | 1,508 [1/2] |
| Nanog Blk Rev[2] | AATGCACACTCCTGAGACCT | NP [2/4] | 810 [2/4] |
| mVenus Rev[3] | GAACTTGTGGCCGTTTA | NP [1/3] | 476 [1/3] |
| YFP-Neo-Fwd[4] | GACGACGGCAACTACAAGAC | | |
| Sox2 Blk Fwd[5] | AGCTCGCAGACCTACATGAA | 710 [5/6] | 1,463 [5/6] |
| Sox2 Blk Rev[6] | TCCTGCAAAGCTCCTACCGTA | NP [6/7] | 803 [6/7] |
| Mid-CFP Fwd[7] | GACGACGGCAACTACAAGAC | NP [5/8] | 680 [5/8] |
| Mid-CFP Rev[8] | GTCTTGTAGTTGCCGTCGTC | | |
| Oct4 Blk Fwd[9] | AGTGGTCCAAGCCTGTAGTC | 905 [9/10] | 1,622 [9/10] |
| Oct4 Blk Rev[10] | GTGACCACTTCCCCATCAGG | NP [10/12] | 950 [10/12] |
| Mid-mCherry Rev[11] | TCAAGTAGTCGGGGATGTCG | NP [9/11] | 692 [9/11] |
| Mid-mCherry Fwd[12] | CGACATCCCCGACTACTTGA | | |

**Table S2. Oligo sequences for primers used in Figure S2.** NP, no product.

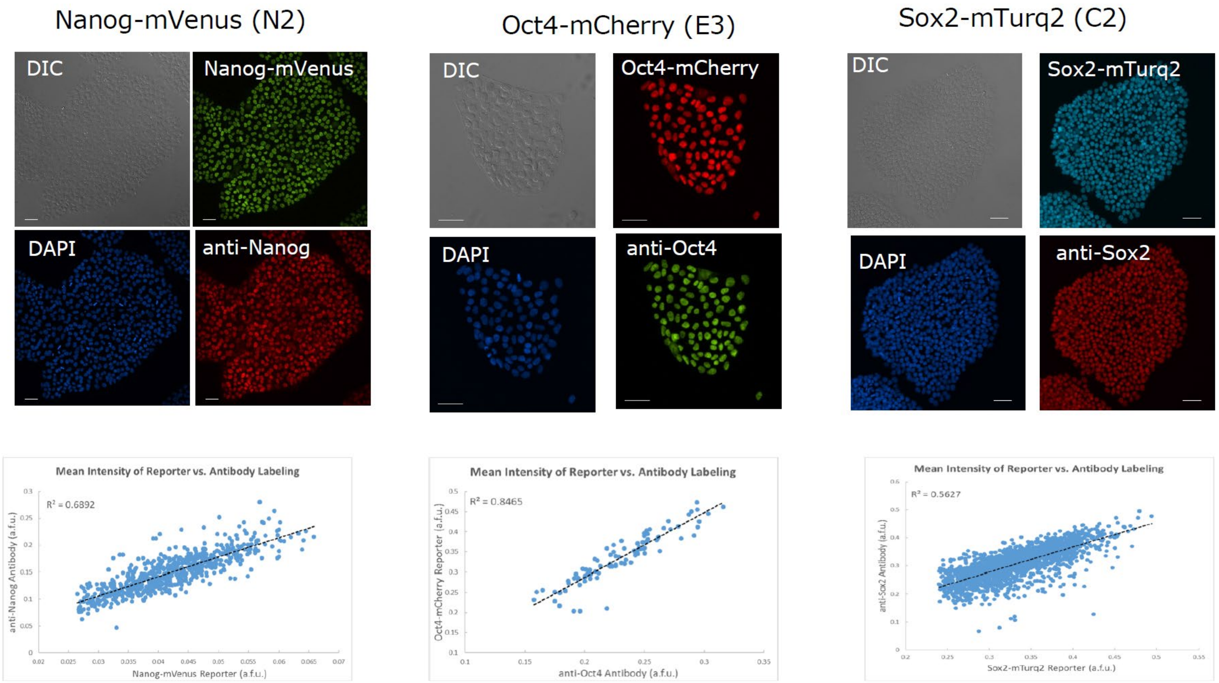


**Figure S3. Comparison of endogenous and reporter protein expression.** Each reporter cell line was fixed and labeled with the corresponding antibody for endogenous protein expression to determine how well the fluorescent protein tag correlated with total protein levels. Mean fluorescence intensity for individual nuclei were segmented using Cell Profiler.

| Genomic variants | OCT4-mCherry | NANOG-mVenus | SOX2-mTurquoise | SOX2-mTurquoise2 |
|---|---|---|---|---|
| Predicted off-target regions | 73 | 55 | 1000+ | 1000+ |
| Predicted off-target regions in genes | 0 | 0 | 0 | 0 |
| SNPs in predicted CRISPR off-target regions | 0 | 0 | 0 | 0 |
| SNPs in high-severity regions | 1 | 0 | 0 | 1 |

**Table S1. Genomic alterations introduced during development of reporter cell lines.**
We performed whole genome sequencing of the parental H9 and reporter cell lines to assess to what extent development of the live-cell reporter cell lines introduced any genomic alterations. We achieved ~30x coverage in each cell line and used GATK[1] to map reads to the H9 reference genome. We then used Strelka[2] to identify 2958 high-confidence SNPs that differed between the parental and derived cell lines. 50 of these SNPs occurred in regulatory regions as determined by H9-specific ATAC-seq[3]. Among those, only 2 SNPs were in high severity regions (i.e., stop-gain or splice site change). None were in high severity regions (i.e., functional regions according to ChromHMM). No SNPs were found in key genes such as *TP53*, *CDKN2A*, *RAS*, or genes frequently mutated in stem cell lines[4]. Deskgen was used to predict the top 20 CRISPR off-targets. High severity regions were predicted off-targets cut off at 1000. One of the variants overlapped *NOTCH2* near a predicted enhancer.

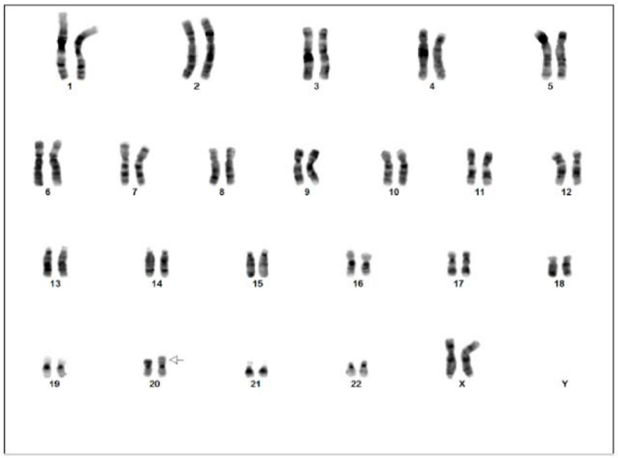

Nanog N2 karyotype

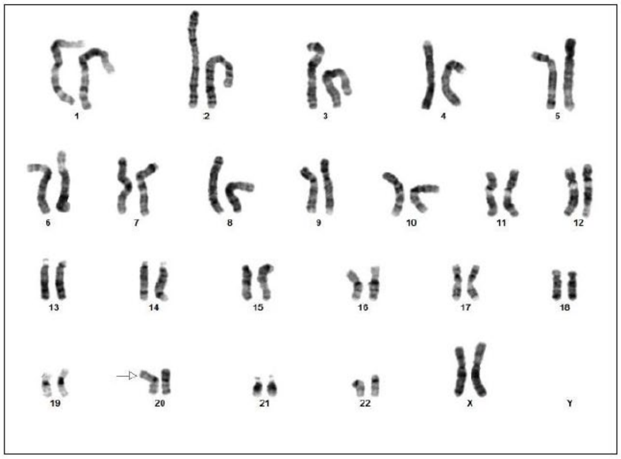

Oct4 E3-10 karyotype

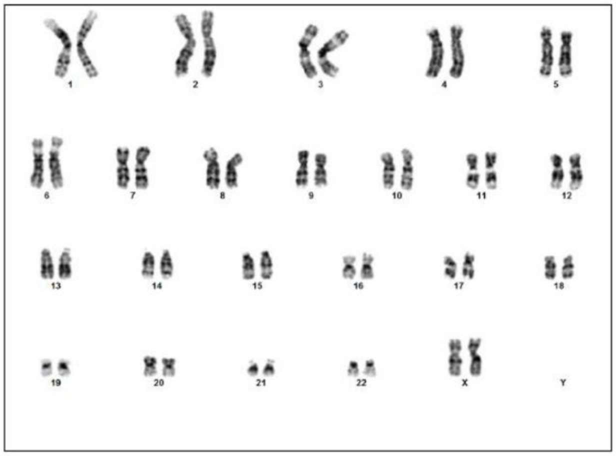

Sox2 C2 karyotype

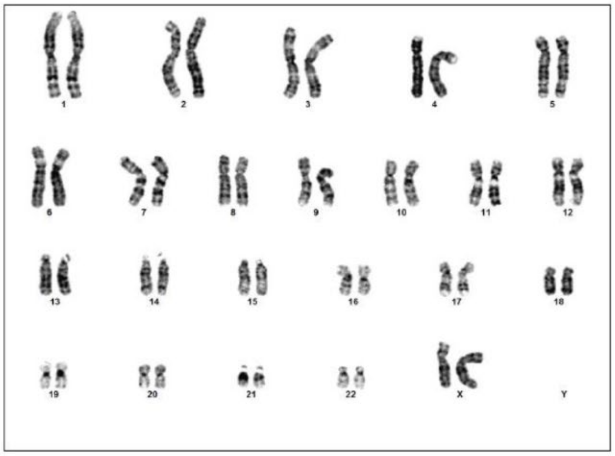

Sox2 C10 karyotype

**Figure S4. Karyotyping analysis of clonal reporter cell lines.**

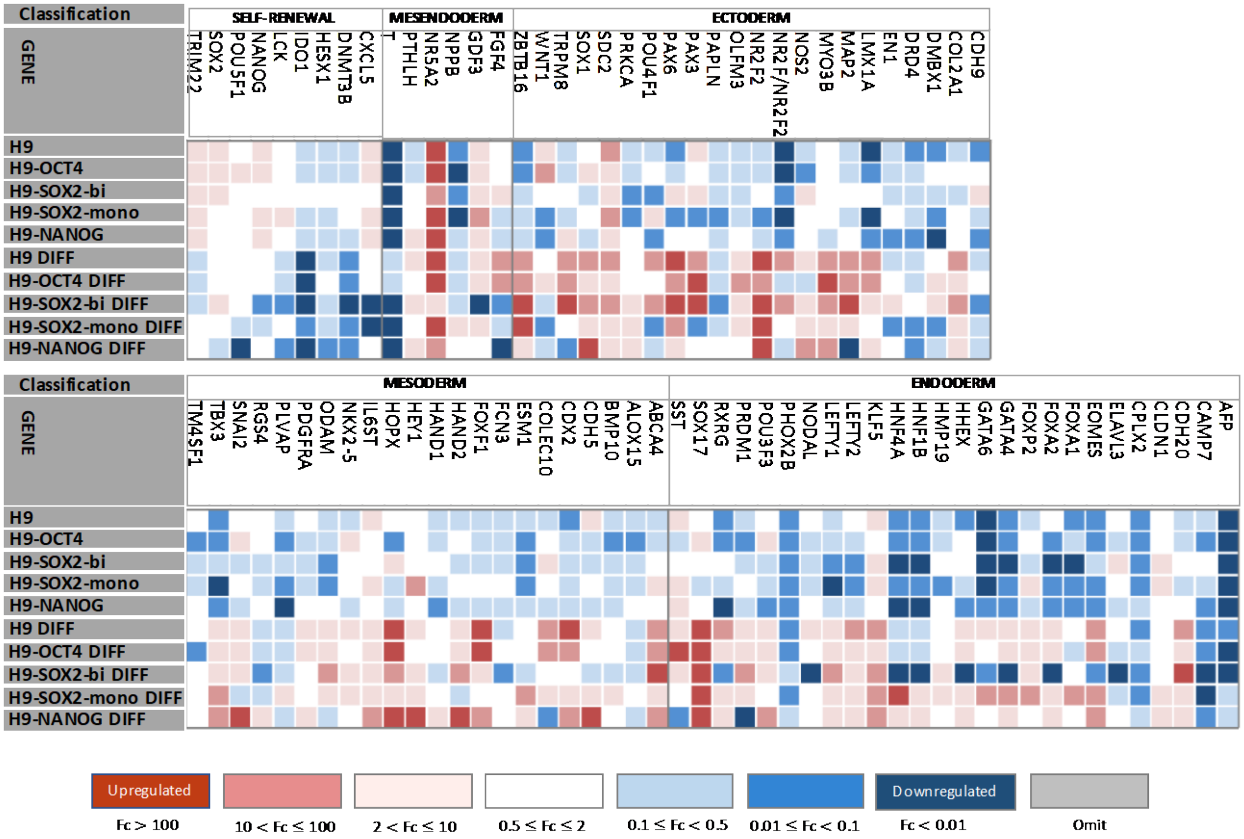


**Figure S5. Transcription profiles of reporter cell lines upon directed differentiation.** To assess that the pluripotency, differentiation potential, and self-renewal capabilities of the reporter cell lines are equivalent to unedited lines, all pluripotent TF reporter cell lines were differentiated to the three germ layers of neuroectoderm, mesoderm, and definitive endoderm then harvested for transcriptomic analysis. The analysis consisted of a qPCR TaqMan hPSC Scorecard Panel85 of 85 gene markers associated with pluripotent, neuroectoderm, mesoderm, and definitive endoderm cell states. All pluripotent factor cell lines had potential for trilineage differentiation and pluripotency.

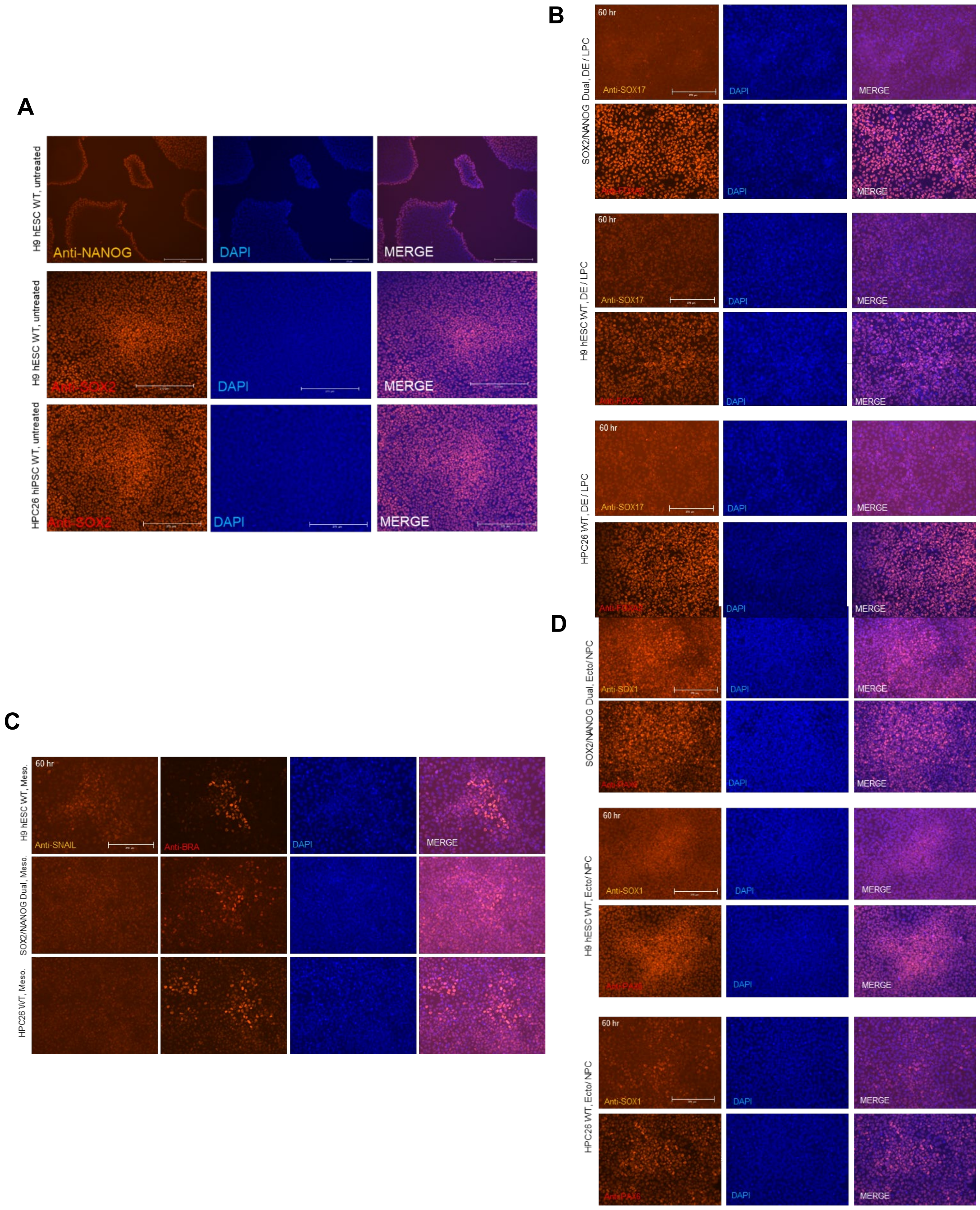


**Fig. S6: Immunofluorescence validation for proper differentiation and unperturbed self-organization.** **A.** IF staining of NANOG and SOX2 in dual reporter, H9 WT, and hiPSC HPC26[5] WT show self-organization and neural rosette formation during unperturbed growth. **B.** IF staining of SOX1 and PAX6 60 hours after ectoderm / NPC differentiation induction in dual reporter, H9 WT, and hiPSC HPC26 WT show positive staining indicative for proper differentiation. **C.** IF staining of SNAIL and BRACHYURY 60 hours after mesoderm / endothelial differentiation induction in dual reporter, H9 WT, and hiPSC HPC26 WT show positive staining indicative for proper differentiation. **C.** IF staining of SOX17 and FOXA2 60 hours after endoderm / LPC differentiation induction in dual reporter, H9 WT, and hiPSC HPC26 WT show positive staining indicative for proper differentiation. Antibody information in Methods section.

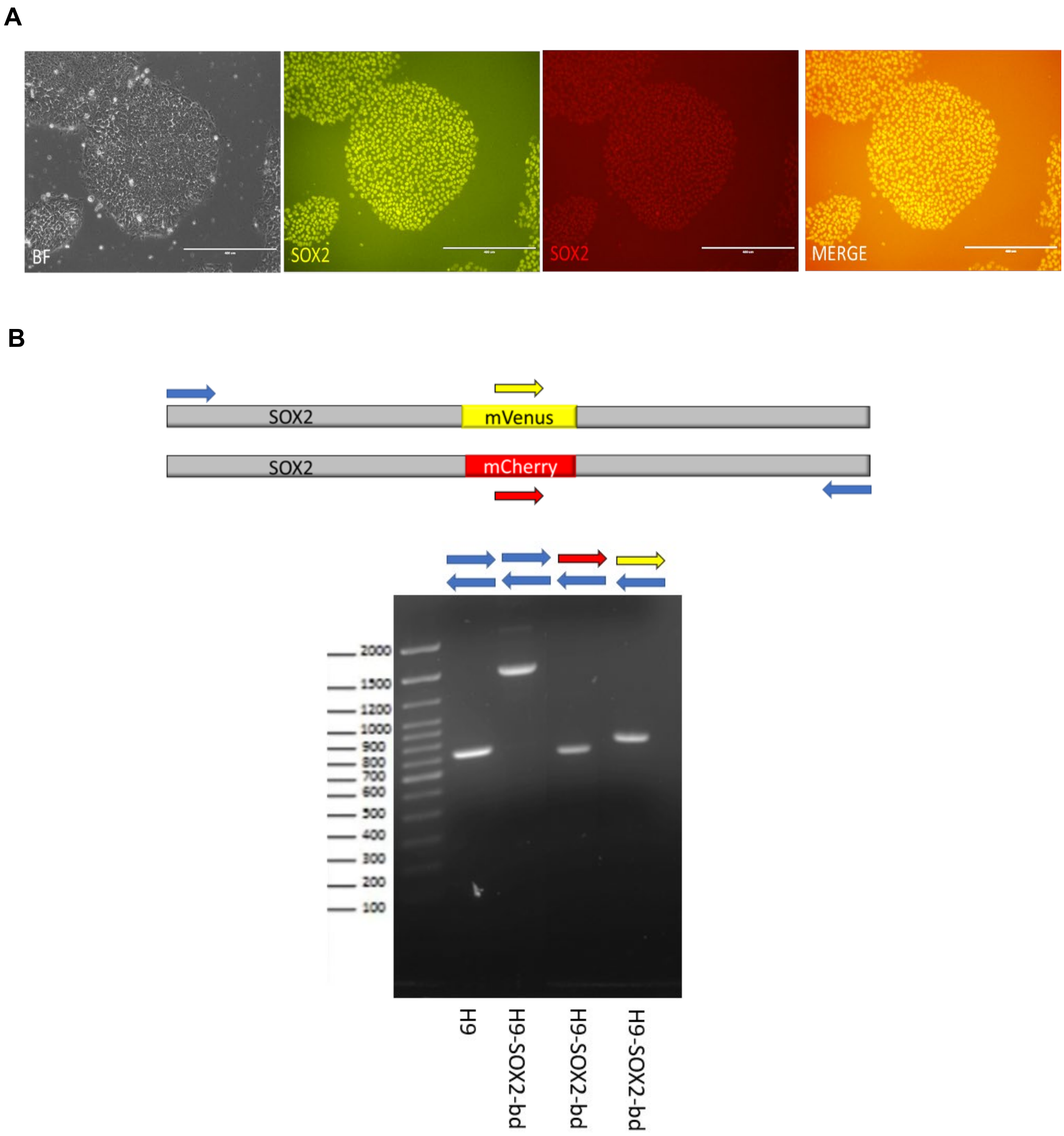


**Fig. S7: SOX2-mVenus/SOX2-mCherry (SOX2 Biallelic Reporter) Cell Line Validations. A.** SOX2-mVenus/SOX2-mCherry biallelic double-tagged line fluorescent microscopy images with from left to right: Brightfield, SOX2-mVenus, SOX2-mCherry, Merge. **B.** Genomic DNA was extracted from H9 SOX2-mVenus/SOX2-mCherry (H9-SOX2-bd) and subjected to PCR with a set of primers flanking the targeted gene. The arrowheads indicate primers used and direction.
